# Cross-Linking of Catalytically Essential Vicinal Thiols at Active Sites of the Cerebral Sodium Transporter Inactivates its Electrogenic Function

**DOI:** 10.1101/2022.10.30.514413

**Authors:** Titilayo Ibironke Ologunagba, Bunmi Olaoluwa Olorundare, Ige Joseph Kade

**Affiliations:** The Federal University of Technology, Akure; Department of Biochemistry, The Federal University of Technology, Akure, Nigeria

**Keywords:** Na^+^/K^+^-ATPase, thiols, cross-linking, diamide, iodoacetamide, dithiothreitol

## Abstract

The inactivation of the electrogenic function of the transmembrane sodium transporter in oxidative stress conditions has been intrinsically linked with the oxidation of its catalytically essential thiols. However, the spatial proximity of these catalytically relevant thiols is yet to be fully elucidated and thus still open. Herein, the influence of a thiol cross-linking [diamide, DA (0.1-2mM)] and a thiol alkylating [iodoacetamide, IA (0.1-5mM)] agent on the activity of the synaptosomal Na^+^/K^+^-ATPase were determined. In addition, the ability of dithiothreitol to either prevent or reverse the inhibition imposed by the thiol modifiers on the enzyme activity was also evaluated. The results showed that the thiol cross-linker inactivates the electrogenic function of the synaptosomal Na^+^/K^+^-ATPase when exposed to the thiols located at either the nucleotide or cation-binding sites. Conversely, irrespective of the exposed active sites, the thiol alkylating agents have no overt effect on the activity of the pump. Furthermore, dithiothreitol markedly prevented but did not reverse the inactivation of the electrogenic pump caused by cross-linking of its critical thiols. Interestingly, both the thiol cross-linker and alkylating agents markedly oxidize dithiothreitol in a time and concentration-dependent fashion. Consequently, within the limit of the present data, it appears that the catalytically relevant thiols of the transmembrane electrogenic pump located at the cationic and nucleotide binding sites, are in close proximity sufficient enough to allow for their cross-linking.

**Highlights:**

- The presence of Na^+^/K^+^-ATPase catalytically important thiols at the nucleotide and cationic sites of the enzyme define its vulnerability to oxidative assault.
- The spatial location of these thiols at vicinal positions at these domains favour the formation of disulphide linkages under oxidative conditions
- The disulphide crosslinking of these thiols culminate in enzyme inactivation
- The inactivation can be prevented but not reversed by exogenous thiol compound

## INTRODUCTION

The central role of the Na^+^/K^+^-ATPase in the normal functioning of the central nervous system has been unequivocally emphasized in literature (Harris *et al*., 2012; Azarias *et al*., 2013; Crambert *et al*., 2000; Zahler *et al*., 1997; Bøttger *et al*., 2011). In fact, its dysfunction has been associated with various oxidative stress-mediated neurodegenerative diseases such as Parkinson’s disease, Alzheimer’s disease, Multiple Sclerosis and several others (Lees, 1991; Liguri *et al*., 1990; Tavalin *et al*., 1997; Chauhan *et al*., 1997; Kumar and Kurup, 2002). Primarily, its critical function involves the active transport of Na^+^ and K^+^ across the plasma membrane, leading to the generation of an electrochemical gradient necessary for driving a wide range of physiological processes (Shrivastava *et al*., 2018; Skou, 1957). Such processes include membrane polarization, glutamate uptake from synaptic junction, regulation of cell volume as well as a number of signalling pathways and consequently cell survival or death (Lees, 1991; Liguri *et al*., 1990; Tavalin *et al*., 1997; Chauhan *et al*., 1997; Kumar and Kurup, 2002).

Therefore, elegant approaches have been directed towards understanding the nature, structure and functions of the Na^+^/K^+^-ATPase (Ohta *et al*., 1986; Lees, 1991; Kaplan 2002). Thus far, efforts have been targeted at elucidating the crystal structure of the enzyme through purification and characterization studies. Reports from such studies have provided extensive information on the various subunits of the pump and have paved way for exploration of the distribution as well as function of the different subunits of this crucial enzyme. (Kaplan, 2002; Morth *et al*., 2007; Shinoda *et al*., 2009). More so, some of these reports have also unveiled the interactions of the enzyme with its substrates, details about its catalytic cycle as well as its regulation (Clausen *et al*., 2017; Toyoshima *et al*., 2011; Shrivastava *et al*., 2018). One of the negative regulators of the enzyme’s function is oxidative stress (Putreshanko *et al*., 2017; Kade *et al*., 2008).

It has been established that one of the critical indices of oxidative stress is the thiol redox status (Kade *et al*., 2008, 2009). Consequently, the susceptibility of the electrogenic sodium pump to oxidative stress is largely due to the presence of catalytically essential sulfhydryl groups at its active site, hence its designation as a sulfhydryl protein (Kaplan, 2002). Therefore, a perturbation of the thiol redox balance poses prodigious threat to the structure and activity of the Na^+^/K^+^-ATPase (Jamme *et al*., 1995; Rauchova *et al*., 1999; Omotayo *et al*., 2015).

In fact, Petrushanko and his co-workers demonstrated that cysteine residues at the actuator domain and nucleotide-binding domains regulate the enzyme’s response to hypoxic conditions (Petrushanko *et al*., 2012, 2017). Therein, it was observed that while Cys 244 essentially regulates the enzyme’s hydrolytic activity, Cys 458 and 459 regulate its signalling function in hypoxic conditions (Petrushanko *et al*., 2012). In consonant with these reports, our group have consistently demonstrated the existence of catalytically critical thiols at the nucleotide and cationic binding sites of the electrogenic sodium pump (Kade *et al*., 2008; Kade, 2012; Kade, *et al*., 2012; Omotayo *et al*., 2011, 2015). Our conclusion was based on the fact that exogenous thiols could either prevent or reverse the inhibition imposed on the transport function of the transmembrane protein by thiol oxidants, such as heavy metals and organochalcogens.

Earlier studies by Kade and coworkers (Kade *et al*., 2012) demonstrated the thiol peroxidase-like activity of organoselenium compounds diphenyl diselenide (DPDS) and dicholesteroyl diselenide (DCDS) on mono and dithiols, as well as sulfhydryl proteins. It was submitted that a lower redox potential of the monothiol, glutathione (GSH) than that of dithiol was responsible for observed differential reactivity with the organochalcogens, wherein dithiothreitol was observed to prevent as well as reverse enzyme inactivation via the thiol peroxidase-like activity of DPDS (Kade et al., 2008). Furthermore, enzyme inactivation by inorganic mercury was significantly prevented in the presence of cysteine. A similar phenomenon was also observed with iron (II), wherein enzyme inactivation by iron (II) was prevented but not reversed by exogenous monothiols, cysteine and glutathione, while dithiothreitol displayed a remarkable potential to prevent and reverse enzyme inactivation. Thus, the prevention or reversal of enzyme inactivation by exogenous thiols is linked to a preferential interaction of the thiol oxidants with exogenous thiols than catalytically relevant thiols of the Na^+^/K^+^-ATPase. Therefore, it is likely that the reductive environment of the exogenous thiols possibly reflects the reductive environment of the critical thiols of the pump at the substrate-binding sites. In fact, higher concentrations of glutathione significantly inhibited the activity of the pump (Kade *et al*., 2012), and this was related to the reductive potential of glutathione being comparatively less than cysteine, another monothiol compound or dithiols, such as dithiothreitol.

Therefore, the differential effects of the thiols [mono or dithiols] on their efficacy to deter or relieve the oxidation of the catalytically essential thiols of the transmembrane transporter may be related in part to the reductive potentials of the different exogenous thiols that were employed. Generally, the dithiols are more potent reductants than monothiols and the former are more efficacious in preventing and relieving inhibition of the pump’s activity imposed by previously exposed thiol oxidants. The inability of the mono thiols to reverse inhibition of pump’s activity by these thiol oxidants may in part suggest that the sulfhydryl groups at the active sites of the cerebral sodium pump is possibly more reductive. This may further suggest the possible existence of multiple thiol groups that are in close proximity at these active sites. However, there is no experimental data to clarify this speculation. Hence the need for this study.

## Materials and Methods

### Chemicals

Adenosinetriphosphate (ATP), diamide, iodoacetamide, dithiothreitol, ouabain, 5’5’-dithio-bis (2-nitrobenzoic) acid (DTNB) were obtained from Sigma (St. Louis, MO). All other chemicals which are of analytical grade were obtained from standard commercial suppliers.

### Animals

Male adult Wistar rats (200–250g) from our own breeding colony were used. Animals were kept in separate animal cages, on a 12-h light: 12-h dark cycle, at a room temperature of 22–24°C, and with free access to food and water. The animals were used according to standard guidelines on the Care and Use of Experimental Animal Resources.

### Preparation of synaptosomal fraction

Rats were decapitated under mild ether anaesthesia and the cerebral tissue (whole brain) was rapidly removed, placed on ice and weighed. The brain was immediately homogenized in cold 100mM Tris–HCl containing 300mM sucrose, pH 7.4 (1/10, w/v) with 10 up-and-down strokes at approximately 1200 rev/min in a Teflon–glass homogenizer. The homogenate was centrifuged for 10 min at 4000g to yield a pellet that was discarded and a low-speed supernatant(S1). The supernatant S1 was then centrifuged at 12000rpm for 10 minutes to yield a membrane and mitochondria-rich synaptosomal fraction as pellet and cytosolic fraction as supernatant. The supernatant was discarded and the pellet was reconstituted in 50mM Tris-HCl, pH 7.4 (1/5, w/v).

### Incubation system for sodium pump assay

Aliquot of the synaptosomal fraction was used for the assay of Na^+^/K^+^-ATPase activity. The reaction mixture contained 3mM MgCl_2_, 125mM NaCl, 20mM KCl and 50mM Tris–HCl, pH 7.4, diamide (final concentrations range of 0.1–2mM), iodoacetamide (final concentrations range of 0.1–5mM), with and without dithiothreitol, cysteine or glutathione (final concentration, 2 mM), and 100–180 mg protein in a final volume of 500 ml. The reaction was initiated by addition of ATP to a final concentration of 3.0mM. Controls were carried out under the same conditions with the addition of 0.1mM ouabain. The reaction mixture was incubated at 37°C for 30 min. At the end of the incubation period, tubes were assayed for sodium pump activity.

### Assay of sodium pump

The reaction system for the assay of the activity of cerebral Na^+^/K^+^-ATPase was essentially the same as described above under the section “incubation system for sodium pump assay”. However, at the end of the incubation time period (30–60 min), the reaction was stopped by addition of 5% trichloroacetic acid. Released inorganic phosphorous (Pi) was measured by the method of Fiske and Subbarow (1925). Na^+^/K^+^-ATPase activity was calculated by the difference between two assays (with and without ouabain). All the experiments were conducted at least three times and similar results were obtained. Protein was measured by the method of Lowry *et al*. (1951), using bovine serum albumin as standard. For all enzyme assays, incubation times and protein concentration were chosen to ensure the linearity of the reactions. All samples were run in duplicate. Controls with the addition of the enzyme preparation after mixing with trichloroaceticacid (TCA) were used to correct for non-enzymatic hydrolysis of substrates. Enzyme activity was expressed as nmol of phosphate (Pi) released min^-1^ mg protein^-1^.

### Oxidation of thiols

The rate of dithiothreitol (DTT) oxidation was determined in the presence of 50mM Tris–HCl, varying pH and at pH 7.4 in the presence of diamide [2mM] and iodoacetamide [2mM]. The rate of thiol oxidation was evaluated by measuring the disappearance of –SH groups. Free –SH groups were determined according to Ellman (1959). Incubation at 37°C was initiated by the addition of DTT (final concentration, 1mM). Aliquots of the reaction mixture (100 ml) were checked (at 1h interval for 3h) for the amount of –SH groups at 412 nm by the addition of the colour reagent 5’5’-dithio-bis (2-nitrobenzoic) acid (DTNB).

### Statistical analysis

Results were analysed by appropriate analysis of variance (ANOVA) and this is indicated in text of results. Duncan’s Multiple Range Test was applied. Differences between groups were considered to be significant when p <0.05.

## Results

### Effect of thiol modifiers on oxidation of thiols-chemical model

The oxidative effect of diamide (DA) and iodoacetamide (IA) on dithiothrietol was investigated. Figure 1 presents the oxidation of DTT [1mM] by DA (panel A) and IA (panel B). A time-dependent oxidation of dithiothreitol by diamide and iodoacetamide was observed.

**Figure 1:**
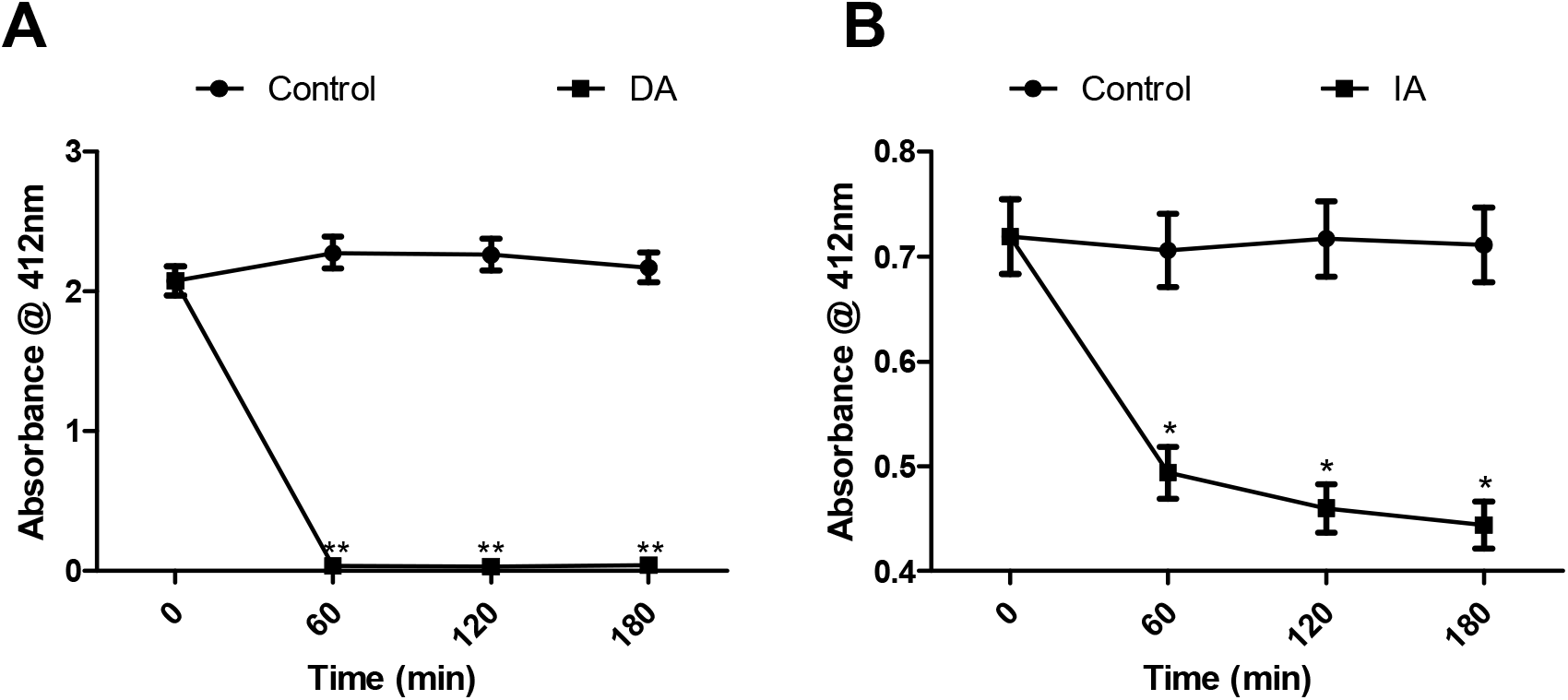
Chemical model of the effect of DA [2mM] (panel A) and IA [2mM] (panel B) on the rate of oxidation of DTT [1mM]. The rate of oxidation was evaluated at the indicated time-points. Data were the means of three independent experiments carried out on different days. Data were presented as mean ± SEM of at least three independent experiments carried out on different days. * indicates significantly lower than control (p < 0.05).

### Effect of thiol modifiers on the activity of the pump

In order to understand the import of the result of the chemical model in biological systems, a simple biological model *in vitro* of this model was carried out using membrane-rich synaptosomal fraction of the cerebral tissue of Wistar albino rats, to evaluate the effect of diamide and iodoacetamide on the activity of Na^+^/K^+^-ATPase. One-way ANOVA analysis of the results showed that the disulphide cross-linking agent, diamide (DA), mediated inhibition of the sodium pump in a concentration-dependent manner, at the nucleotide-binding site (Figure 2, panel A). However, iodoacetamide had no inhibitory effect on the activity of the electrogenic pump (panel B).

**Figure 2:**
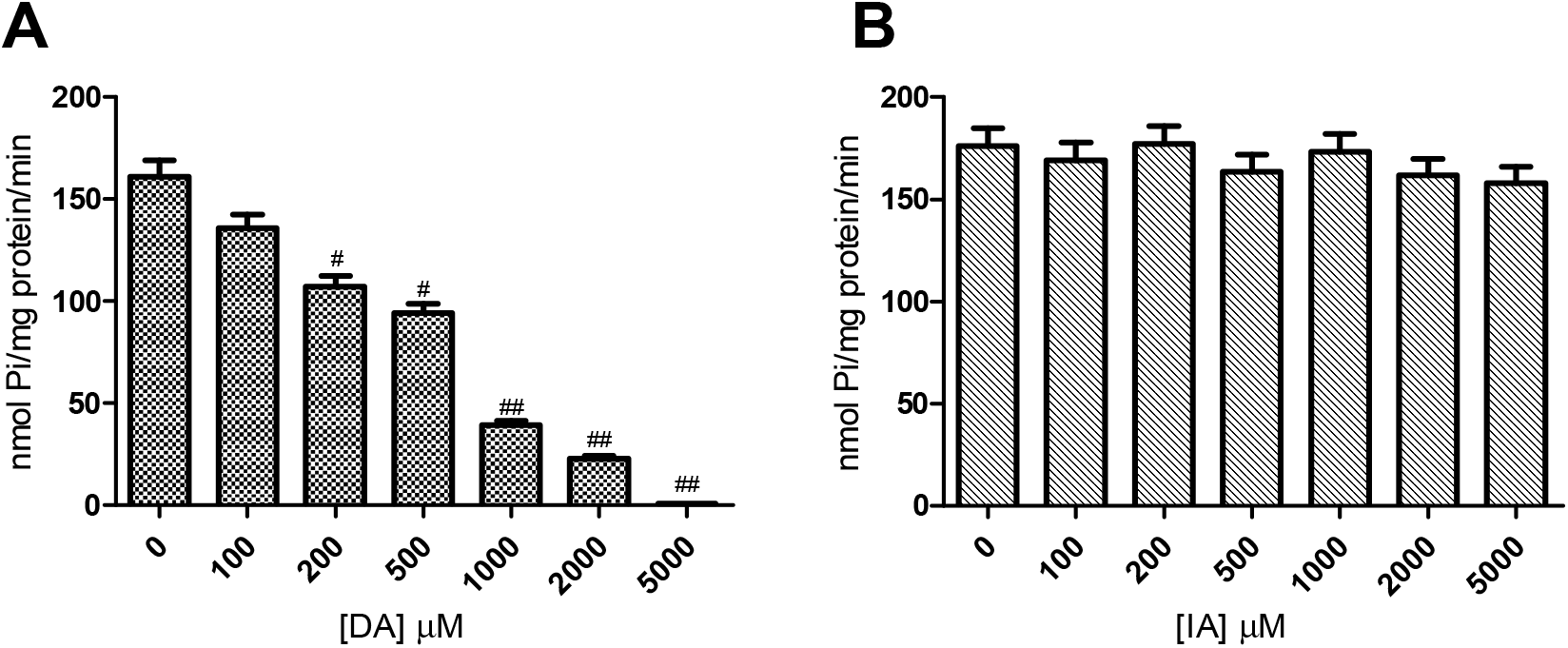
Effect of varying concentrations of disulphide-crosslinking agent, diamide (panel A), and iodoacetamide (panel B) on the activity of the Na^+^/K^+^-ATPase at the nucleotide-binding site. Data were presented as mean ± SEM of at least three independent experiments carried out on different days. # indicates significantly lower than control (p < 0.05).

### Effect of exogenous thiol, dithiothreitol, on DA-mediated enzyme inactivation

Diamide is a thiol-specific disulphide crosslinking agent, and is therefore speculated to interact with the catalytically relevant thiols of this sulfhydryl protein. Hence, the effect of pre- and post-incubation with exogenous thiol, dithiothreitol, on the inhibition mediated by diamide was investigated. As presented in Figure 3, two-way ANOVA analysis of the results revealed that preincubation with DTT (panel A), significantly prevented the inhibition mediated by diamide on the activity of the pump (p < 0.05). In contrast, post-incubation with DTT did not reverse DA-mediated enzyme inactivation (panel B).

**Figure 3:**
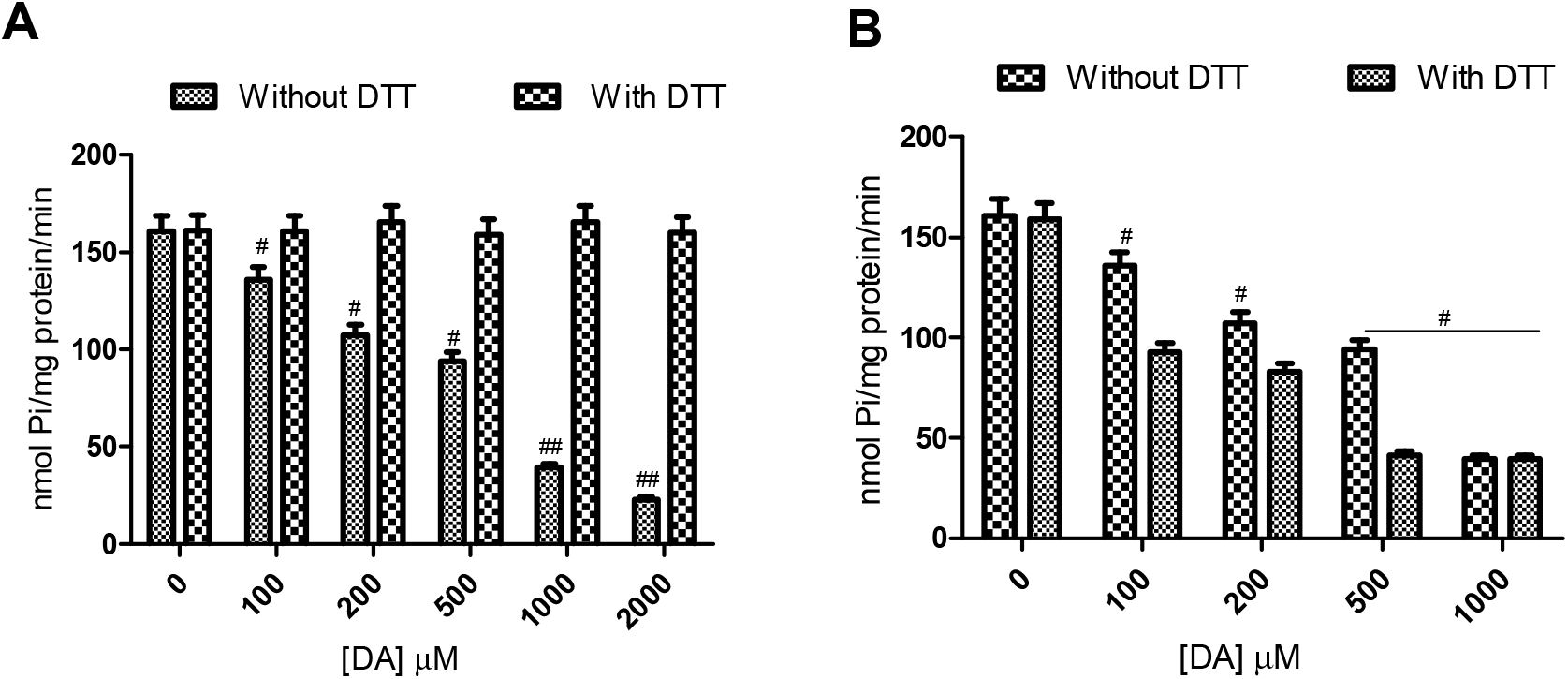
Effect of pre-incubation (panel A) and post-incubation (panel B) with 2mM DTT on diamide-induced inhibition of the Na^+^/K^+^-ATPase at the nucleotide-binding site. Data were presented as mean ± SEM of at least three independent experiments carried out on different days. # indicates significantly lower than control (p < 0.05) while * indicates significantly higher than corresponding concentration without DTT.

### Effect of diamide and iodoacetamide on Na^+^/K^+^-ATPase activity at the cation-binding sites

Furthermore, based on speculations of thiol groups at the cation-binding sites of this protein, the effect of the thiol modifying agents on the activity of the sodium pump when the cation-binding sites were exposed was investigated. Figure 4 showed (one-way ANOVA analysis) that diamide inhibited the activity of the pump significantly (p < 0.05), in a concentration dependent fashion, when Na^+^, K^+^ and Mg^2+^ were excluded from the pre-incubating medium (panels A, C and E respectively). However, iodoacetamide had no significant effect on the activity of the pump at all three cationic sites (panels B, D and F respectively in the same order as for diamide).

**Figure 4:**
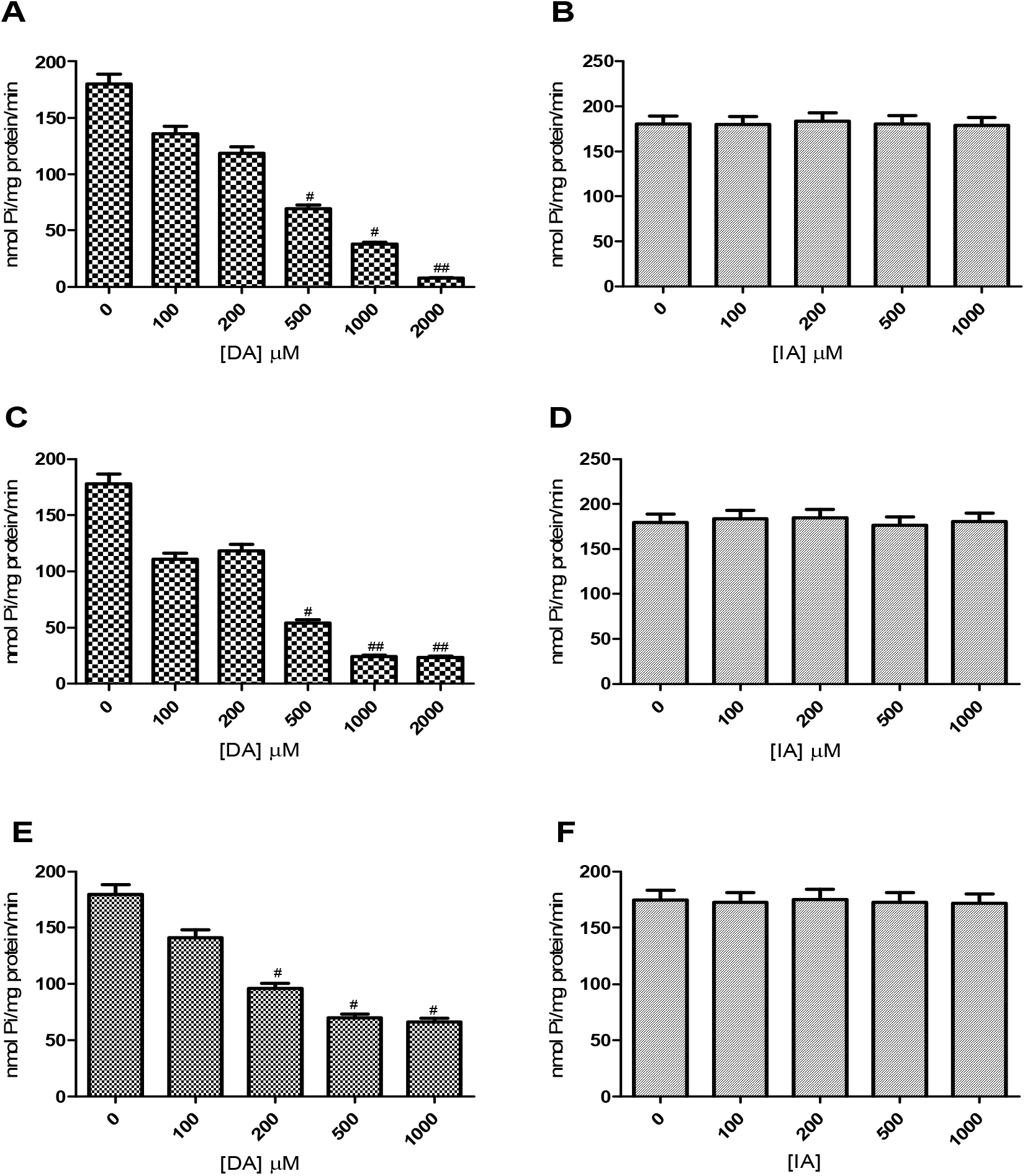
Effect of varying concentrations of diamide and iodoacetamide on the activity of the sodium pump at the Na^+^ (A and B respectively), K^+^ (C and D respectively) sites and upon exclusion of Mg^2+^ from the preincubation medium (E and F respectively). Data were presented as mean ± SEM of at least three independent experiments carried out on different days. # indicates significantly lower than control (p < 0.05).

### Effect of exogenous thiol, DTT, on DA-mediated enzyme inactivation at the cation-binding sites

Similar to the investigation of the nucleotide binding site, we examined the effect of pre- and post-incubation with DTT on the inactivation mediated by diamide on the activity of the pump when each of the cations were excluded from the preincubating medium. Two-way ANOVA analysis of the result is as presented in Figure 5, wherein preincubation with DTT significantly prevented the inhibition induced by diamide in the absence of each of the cations Na^+^ (panel A), K^+^ (panel C) and Mg^2+^ (panel E) from the preincubating medium (p < 0.05). However, post-incubation with DTT did not reverse the inhibition by diamide for all three cation-binding sites (panels B, D and F respectively).

**Figure 5:**
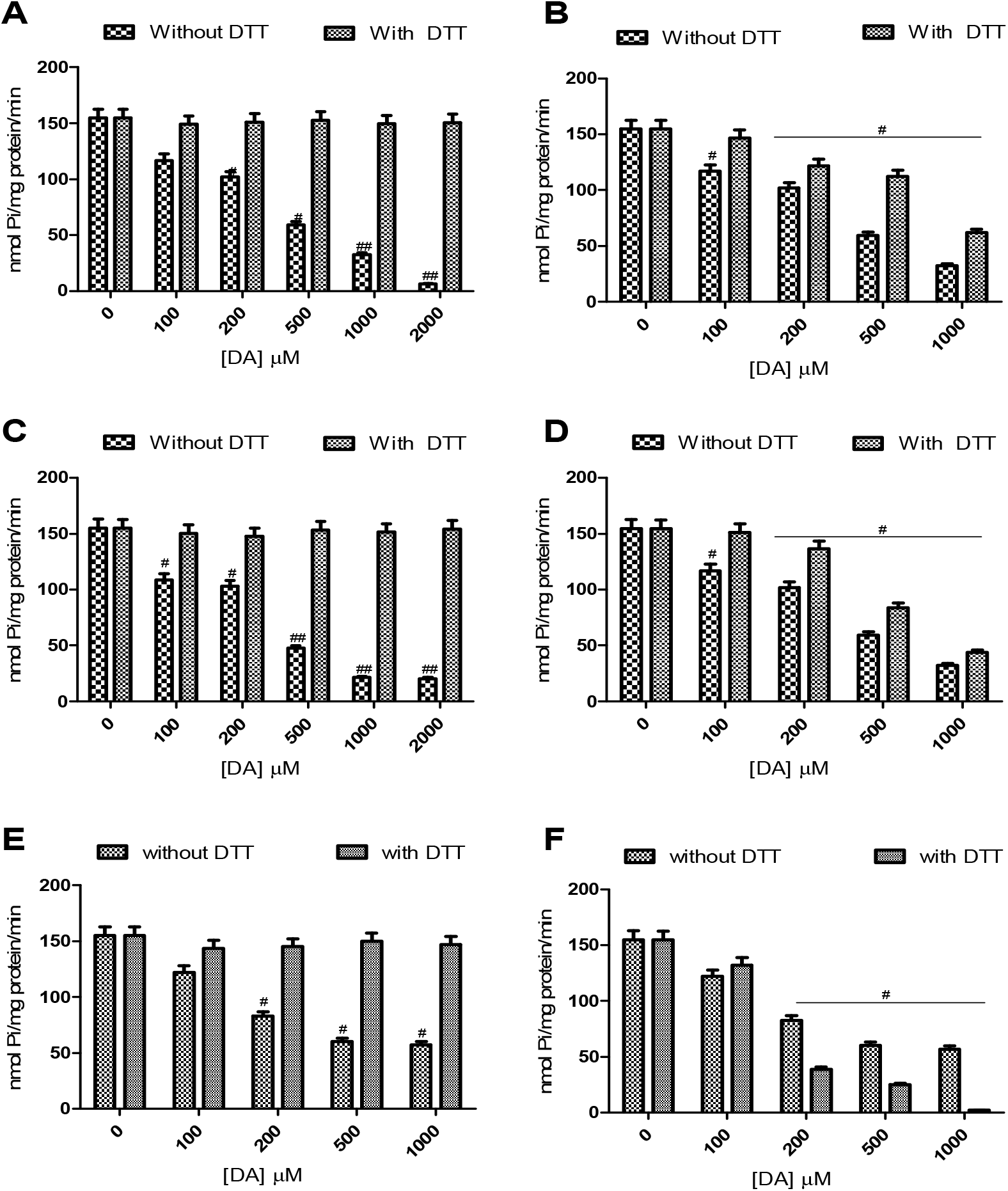
Effect of preincubation with 2mM DTT on diamide-induced inhibition of the Na^+^/K^+^-ATPase when Na^+^ (panel A), K^+^ (panel C), and Mg^2+^ (panel E) respectively were excluded from the preincubating medium, and the effect of post-incubation with DTT on diamide-induced inhibition upon exclusion of each of the respective cations from the preincubating medium (panels B, D and F respectively). Data were presented as mean ± SEM of at least three independent experiments carried out on different days. # indicates significantly lower than control (p < 0.05).

## Discussion

The oxidation of the critical thiols of the Na^+^/K^+^-ATPase has been central to the discourse about inhibition of the enzyme’s activity under different physiological conditions (Nogueira *et al*., 2004; Kade *et al*., 2008; Omotayo *et al*., 2011; 2015; Petrushanko *et al*., 2012; 2017). In these reports, enzyme inhibition was found to be prevented in the presence of exogenous thiols (dithiols and monothiols). Some of these reports employed a chemical model of the oxidation of mono- and di-thiols to investigate the molecular events involved in the oxidation of this sulfhydryl protein by thiol oxidizing agents (Kade *et al*., 2008; Omotayo *et al*., 2011; 2015; Petrushanko *et al*., 2012; 2017). In the present study, a disulphide crosslinking agent, diamide (DA) as well as a thiol alkylating agent, iodoacetamide (IA), were employed in chemical and simple biological models to verify the possible presence of vicinal thiols at the substrate-binding sites of the Na^+^/K^+^-ATPase. Using a chemical model, the oxidation of dithiothreitol by diamide and iodoacetamide showed that both thiol modifiers mediated oxidation of thiols in a time-dependent fashion (Figure 1). Specifically, the gradual and time-dependent manner of DA-mediated oxidation of 2mM DTT (data not shown) in contrast with the drastic and near-complete oxidation of 1mM DTT suggests a stoichiometric dimension of diamide thiol oxidation. In fact, it has been reported that the reaction of DA with free thiols proceeds by the abstraction of hydrogen atom and its consequent reduction to an unstable sulfenylhydrazine intermediate which is further reduced by oxidation of another thiol group to generate the hydrazine product and a disulphide (Leichert *et al*., 2003). On the other hand, iodoacetamide modifies thiol groups by bimolecular nucleophilic substitution reactions to form carbamidomethylated cysteines-a second order reaction that is dependent on the concentration of both the nucleophile (S_) and iodoacetamide (the substrate), as well as pH of the solvent (Hill *et al*., 2009; Wong and Liebler, 2008; Rogers *et al*., 2006). It is apparent that the reaction of diamide with thiols is essentially completed in the presence of two thiol groups (Scheme 1). This may possibly explain the near-complete oxidation of other mono (data not shown) and di-thiols by diamide used in the present experiment and further buttresses the established disulphide-crosslinking property of DA. This chemical model can be used to interpret the result obtained from the simple biological model *in vitro*.

**Scheme 1:**
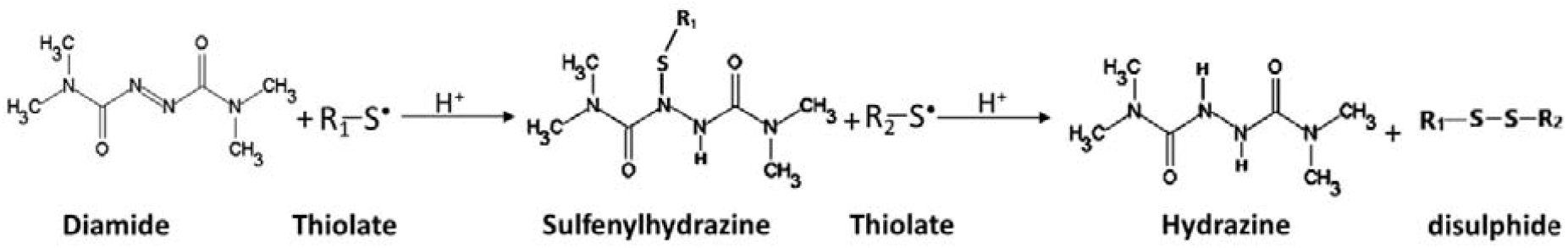
Scheme showing the reaction of diamide with free thiols.

The Na^+^/K^+^-ATPase is a sulfhydryl protein having catalytically relevant thiol groups at its substrate-binding sites (Kurella *et al*., 1999; Kade *et al*., 2008, 2009; Omotayo *et al*., 2011; 2015). The inhibition of the Na^+^/K^+^-ATPase by DA, at the ATP-binding site (Figure 2, panel A), corroborates previous submissions about the presence of catalytically essential thiols at this site. However, the inability of iodoacetamide to inhibit the electrogenic function of this pump is a puzzle (panel B), since the chemical model, as well as other reports (Hill *et al*., 2009; Wong and Liebler, 2008), have demonstrated its thiol oxidizing potential. Therefore, to ascertain the involvement of the critical thiols of the nucleotide-binding site in DA-mediated inhibition, exogenous thiol, DTT, was introduced into the preincubating medium. The observed ability of DTT to protect enzyme from inhibition mediated by DA validates the modification of the catalytically relevant thiols of this pump as a primary mechanism involved in DA-mediated enzyme inactivation (Figure 3, panel A). However, post-incubation with DTT did not recover enzyme activity following DA-mediated inactivation (panel B). This phenomenon is at variance with what was observed with another thiol oxidant, diphenyl diselenide (Kade *et al*., 2012) where post-incubation with DTT reversed enzyme inactivation. A number of speculations could be attributed to this phenomenon.

Firstly, evidence from literature suggests that the interaction of DA with thiols yields disulphide and a hydrazine product, as illustrated in Scheme 1, and this is in agreement with the observed pattern of DA-mediated thiol oxidation in the chemical model. Basically, the presence of thiol groups in close proximity at the enzyme’s active sites provides a reductive environment that is sufficiently mimics the reductive environment of the dual thiol groups of DTT which favours the reduction of DA to its hydrazine product. This is likely accompanied by formation of disulphide linkages between these catalytically relevant thiols, and consequent enzyme activation. Therefore, preincubation with DTT supplies a more easily accessible reductive alternative for DA, thus sparing the vicinal catalytically relevant thiols of the pump and preventing consequent enzyme inactivation. In addition, the fact that the formation of disulphide is intricately intertwined with a protein’s tertiary/quartenary structure (Leichert *et al*., 2003) may likely explain the inability of DTT to reverse enzyme inactivation by DA upon post incubation with DTT. Particularly, the formation of disulphide linkages other than the protein’s native disulphide bonds could significantly modify its native conformation. This likely induces conformational alteration of the pump’s tertiary/quaternary structure due to formation of non-native disulphide linkages, which may account for the observed sustained enzyme inactivation.

More so, other reports have implied the presence of vicinal thiols at the catalytic sites of this enzyme as well as another sulfhydryl protein, as δ-ALA-D (Kade *et al*., 2008, 2009; Puntel *et al*., 2010). In these reports, these catalytically relevant thiols were the primary targets of a thiol-oxidizing organoselenium compound, diphenyl diselenide. It was submitted that the proximity of these thiols in the tertiary structure of δ-ALA-D was critical in order to favour the interaction of diphenyl diselenide with these enzymes (Nogueira and Rocha, 2010). Although, this speculation was projected for the Na^+^/K^+^-ATPase, there is no experimental evidence to validate this claim as established in the present study.

Thus, an introduction of DTT (post incubation) into the reaction system is possibly insufficient to reverse such conformational alteration. This could also be due to the fact that the formation of the product of diamide reduction, hydrazine, cannot be reversed by DTT.

The introduction of DTT, in the absence of other electron acceptors, is apparently insufficient to regenerate the N=N double bond of diamide, hence the reaction is not likely reversible.

Furthermore, the α-subunit of the Na^+^/K^+^-ATPase has 23 evolutionarily conserved cysteine residues. Fifteen of these are located in the cytosolic loop forming the ATP-binding site (Kurella *et al*., 1999). Under conditions of low ATP-supply and mild oxidative stress, it has been earlier reported that three of these cysteine residues, Cys-454, -458, and -459, undergo regulatory S-glutathionylation (which is reversible) and results in an increase in negative charge within the ATP-binding socket (Putreshanko *et al*., 2017). Taken together, it is clear from the present study that the ATP-binding site of this crucial enzyme is laced with vicinal sulfhydryl groups that contribute to high reductive potential at this site.

Furthermore, the binding of Mg^2+^ to ATP and formation of the MgATP^2-^ is essential for shielding it from the electronegative intracellular environment and supports the formation of the phosphate groups in the nucleotide. Therefore, it is logical that the exclusion of Mg^2+^ from the reaction medium could induce a structural deformity in the ATP molecule, prevent its binding at the nucleotide-binding site and, hence, favour enzyme inhibition by DA (Figure 4, panel E). The phenomenal inability of IA to inactivate the pump’s electrogenic function (panel F) in this condition further corroborates the fact that a typical feature of iodoacetamide may be responsible for this observation. DA has been reported to rapidly cross biological membranes (Toledano *et al*., 2003; Pócsi *et al*., 2005), being lipophilic, however, iodoacetamide is more hydrophilic. Furthermore, iodoacetamide has been used in experiments involving peptide mapping to prevent the formation of disulphide bonds by the protein (Hill *et al*., 2009). It is not clear whether the prevention of disulphide bond formation by iodoacetamide accounts for its inability to inactivate the electrogenic pump. Further study on this phenomenon may hold insightful details about the microenvironment and location of the critical thiols of the pump. The observation of similar trend following pre and post incubation with DTT (Figure 5, panels E and F respectively) further establishes the speculations earlier described for DA-mediated enzyme inactivation at the nucleotide-binding site.

Moreover, earlier study in our laboratory reported that the cationic site of this enzyme contains sulfhydryl groups that are susceptible to oxidation by thiol oxidizing agents (Omotayo *et al*., 2011). In the present study, diamide inhibited enzyme activity at these cationic sites, buttressing earlier submission (Figure 4, panels A and C respectively). Similar to discussion in the previous section on the ATP-binding site, the inhibition mediated by diamide may involve the formation of disulphide bonds between vicinal thiols at this site. More so, preincubation with DTT protected the enzyme from diamide-mediated inactivation (Figure 5, panels A and C), suggesting the preferential formation of disulphide bond between the thiols of dithiothreitol, and consequently sparing the critical thiols at the cationic sites of the pump. This is logical since we have earlier demonstrated in the chemical model that the reaction of 2mM DTT with 2mM DA displayed a stoichiometric balance that sufficiently interacts over the course of three hours, hence the thirty-minute incubation period for the enzyme assay falls within the time-course of this reaction. The implication of this is that the vicinal and catalytically relevant thiols at the cationic sites of this crucial electrogenic transporter are spared of DA-mediated oxidation, consequently preventing enzyme inactivation in the simple biological model. This explanation also applies to the nucleotide-binding site. However, inability to recover enzyme activity by post-incubation with DTT at the Na^+^ and the K^+^ sites (Figure 5, panels B and D) is likely based on a similar principle as discussed for the nucleotide binding site. Summarily, Scheme 2 suggest the possible events that may characterize enzyme inhibition by diamide and protection by DTT.

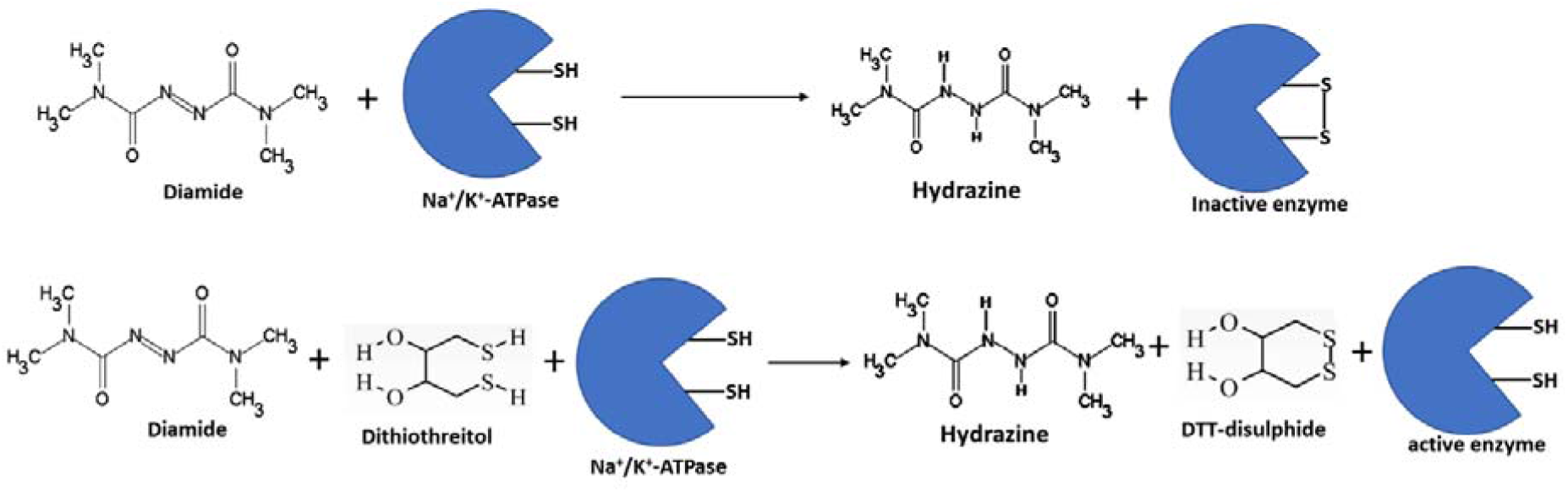

From our findings, both the nucleotide and cationic sites of the cerebral transmembrane electrogenic pump contain vicinal thiols that are susceptible to forming disulphide bonds under oxidative conditions. This leads to enzyme inactivation and possible protein misfolding leading to several lethal consequences, particularly in the brain. More so, since neurodegenerative disease are generally characterized by sustained oxidative stress, the oxidation these vicinal and critical thiols of the Na^+^/K^+^-ATPase may be intricately involved in their pathology.

## Acknowledgement

TIO and IJK acknowledge the financial support of DBT-India and TWAS. The authors also acknowledged the financial support of CAPES, FINEP, FAPERGS, PRONEX and CNPq, FINEP research grant Rede Instituto Brasileiro de Neurociencia (IBN-Net) # 01.06.0842-00 and the INCT for excitotoxicity and Neuroprotection-CNPq and CNPq-ProAfrica grant awarded to IJK and JBTR.

## Notes

### Competing Interest Statement

The authors have declared no competing interest.

## REFERENCES

Azarias, G., Kruusmägi, M., Connor, S., Akkuratov, E. E., Liu, X.-L., Lyons, D., Brismar, H., Broberger, C., Aperia, A. (2013). A specific and essential role for Na,K-ATPase α3 in neurons co-expressing α1 and α3. J. Biol. Chem., 288: 2734–43.

Bavaresco C., Calcagnotto T., Tagliari B, Delwing D., Lamers M. L., Wannmacher C. M., Wajner M., Wyse A. T. (2003). Brain Na,K-ATPase inhibition induced by arginine administration is prevented by vitamins E and C. Neurochem. Res., 28:825–829

Bøttger, P., Tracz, Z., Heuck, A., Nissen, P., Romero-Ramos, M., Lykke-Hartmann, K. (2011). Distribution of Na/K-ATPase alpha 3 isoform, a sodium-potassium P-type pump associated with rapid-onset of dystonia parkinsonism (RDP) in the adult mouse brain. J. Comp. Neurol, 519: 376–404.

Carfagna M. A., Ponsler G. D., Muhoberac B. B. (1996). Inhibition of ATPase activity in rat synaptic plasma membranes by simultaneous exposure to metals. Chem. Biol. Interact, 100:53–65

Chauhan N. B., Lee J. M., Siegel G. J. (1997). Na,K-ATPase mRNA levels and plaque load in Alzheimer’s disease. J. Mol. Neurosci, 9:151–66

Clausen M. V., Hilbers F., Poulsen H. (2017). The Structure and Function of the Na,K-ATPase Isoforms in Health and Disease. Front. Physiol., 8: 371.

Crambert, G., Hasler, U., Beggah, A. T., Yu, C., Modyanov, N. N., Horisberger, J. D., Lelièvre, L., Geering, K. (2000). Transport and pharmacological properties of nine different human Na, K-ATPase isozymes. J. Biol. Chem., 275: 1976–86.

De Assis D. R, Ribeiro C. A, Rosa R. B et al. (2003). Evidence that antioxidants prevent the inhibition of Na,K-ATPase activity induced by octanoic acid in rat cerebral cortex in vitro. Neurochem. Res., 28:1255–1263

Ellman, G. L (1959). Tissue sulfhydryl groups. Arch. Biochem. Biophys, 82: 70–77

Fiske, C. H., Subbarow, Y. J (1925). The colorimetric determination of phosphorus. J. Biol. Chem., 66: 375–381.

Harris, J. J., Jolivet, R., Attwell, D. (2012). Synaptic energy use and supply. Neuron, 75: 762–777.

Hill B. G., Reily C., Oh J.-Y., Johnson M. S., Landar A. (2009). Methods for the determination and quantification of the reactive thiol proteome. Free Radic. Biol. Med., 47(6): 675–683

Jamme I., Petit E., Divoux D., Gerbi A., Maixent J. M., Nouvelot A. (1995) Modulation of mouse cerebral Na+,K+-ATPase activity by oxygen free radicals. Neuroreport, 7(1): 333–337 8742483.

Kade I. J, Paixão M. W, Rodrigues O. E, Barbosa N. B, Braga A. L, Avila D. S, Nogueira C. W, Rocha J. B. (2008). Comparative studies on dicholesteroyl diselenide and diphenyl diselenide as antioxidant agents and their effect on the activities of Na+/K+-ATPase and delta-aminolevulinic acid dehydratase in the rat brain. Neurochem. Res., 33:167–178

Kade I. J., Paixão M. W., Rodrigues O. E., Ibukun E. O., Braga A. L., Zeni G., Nogueira C. W., Rocha J. B. T. (2009). Toxicol In Vitro, 23: 14

Kaplan, J. H. (2002). Biochemistry of Na,K-ATPase. Annu. Rev. Biochem., 71: 511–535

Kosower, N. S., Kosower, E. M. (1995). Diamide: an oxidant probe for thiols. Meth. Enzymol., 251, 123–133.

Kourie J. I. (1998). Interaction of reactive oxygen species with ion transport mechanisms. Am J Physiol, 275:1–24.

Kumar, A. R, Kurup, P. A. (2002). Inhibition of membrane Na+-K+ ATPase activity: a common pathway in central nervous system disorders. J Assoc Physicians India, 50:400–6.

Kurella E. G., Tyulina O. V., Boldyrev A. A. (1999) Oxidative resistance of Na/K-ATPase. Cell. Mol. Neurobiol, 19: 133–140

Lees, G. J. (1991) Inhibition of sodium–potassium-ATPase: a potentially ubiquitous mechanism contributing to central nervous system neuropathology. Brain Res. Rev., 16 (3): 283–300.

Leichert, L. I. O., Scharf, C., Hecker, M. (2003). Global Characterization of Disulfide Stress in Bacillus subtilis. J. Bacteriol., 185(6): 1967–1975

Liguri G., Taddei N., Nassi P., Latorraca S., Nediani C., Sorbi S. (1990). Changes in Na+, K+-ATPase, Ca2+-ATPase and some soluble enzymes related to energy metabolism in brains of patients with Alzheimer’s disease. Neurosci. Lett., 112(2–3): 338–342.

Lowry, O. H., Rosenbrough, N. J., Farr, A. L., Randall, R. J. (1951) Protein measurement with Folin-phenol reagent. J. Biol. Chem., 193: 265–275

Morth J. P., Pedersen B. P., Toustrup-Jensen M. S., Sørensen T. L-M., Petersen J., Andersen J. P., Vilsen B., Nissen P. (2007). Crystal structure of the sodium–potassium pump. Nature, 450: 1043–1049.

Nogueira C. W., Rocha J. B. T. (2010). Diphenyl diselenide: a Janus-faced compound. J Braz Chem Soc, 21:2055–2071.

Nogueira C. W., Zeni G., Rocha J. B. T. (2004) Organoselenium and organotellurium compounds: toxicology and pharmacology. Chem. Rev, 104:6255–6285

Ohta T., Nagano K., Yoshida M. (1986). The active site structure of Na+/K+-transporting ATPase: Location of the 5’-(p-fluorosulfonyl)benzoyladenosine binding site and soluble peptides released by trypsin. Proc. Natl. Acad. Sci. U. S. A., Vol. 83: 2071–2075.

Omotayo T. I., Akinyemi G. S., Omololu P. A., Ajayi B. O., Akindahunsi A. A., Rocha J. B. T., Kade I. J. (2015). Possible involvement of membrane lipids peroxidation and oxidation of catalytically essential thiols of the cerebral transmembrane sodium pump as component mechanisms of iron-mediated oxidative stress-linked dysfunction of the pump’s activity. Redox Biol., 4: 234–241.

Omotayo, T. I., Rocha J. B. T., Ibukun E. O., Kade I. J. (2011). Inorganic mercury interacts with thiols at the nucleotide and cationic binding sites of the ouabain-sensitive cerebral electrogenic sodium pump. Neurochem. Int., 58: 776–784

Petrushanko I. Y., Mitkevich V. A., Lakunina V. A., Anashkina A. A., Spirin P. V., Rubtsov P. M., Prassolov V. S., Bogdanov N. B., Hänggi P., Fuller W., Makarov A. A., Bogdanov A. (2017). Cysteine residues 244 and 458–459 within the catalytic subunit of Na,KATPase control the enzyme’s hydrolytic and signaling function under hypoxic conditions. Redox Biol., 13: 310–319

Petrushanko I. Y., Yakushev S., Mitkevich V. A., Kamanina Y. V., Ziganshin R. H., Meng X., Anashkina A. A., Makhro A., Lopina O. D., Gassmann M., Makarov A. A., Bogdanova A. (2012). S-glutathionylation of the Na,K-ATPase catalytic alpha subunit is a determinant of the enzyme redox sensitivity. J. Biol. Chem., 287: 32195–32205

Pócsi I., Miskei M., Karányi Z., Emri T., Ayoubi P., Pusztahelyi T., Balla G., Prade R. A. (2005). Comparison of gene expression signatures of diamide, H_2_O_2_ and menadione exposed Aspergillus nidulans cultures – linking genome-wide transcriptional changes to cellular physiology. BMC Genom., 6:182

Puntel R. L., Roos D. H., Folmer V., Nogueira C. W., Galina A., Aschner M., Rocha J. B. T. (2010). Mitochondrial dysfunction induced by different organochalcogens is mediated by thiol oxidation and is not dependent of the classical mitochondrial permeability transition pore opening. Toxicol. Sci., 117(1): 133–43.

Rauchová H., Drahota Z., Koudelová J. (1999). The role of membrane fluidity changes and thiobarbituric acid-reactive substances production in the inhibition of cerebral cortex Na+/K+-ATPase activity. Physiol. Res., 48(1): 73–78

Rogers L. K., Leinweber B. L., Smith C. V. (2006). Detection of reversible protein thiol modifications in tissues. Anal. Biochem., 358:171–184.

Schmidt M. M., Dringen R. (2009). Differential effects of iodoacetamide and iodoacetate on glycolysis and glutathione metabolism of cultured astrocytes. Front. Neuroenergetics., 1(1): 1–10

Shinoda T., Ogawa H., Cornelius F., Toyoshima C. (2009). Crystal structure of the sodium– potassium pump at 2.4A° resolution. Nature Lett., 459: 446–451.

Shrivastava A. N., Triller A., Melki R. (2018). Cell Biology and Dynamics of Neuronal Na+/K+-ATPase in Health and Diseases. Neuropharmacology, 169:107461.

Skou, J. C. (1957). The influence of some cations on an adenosine triphosphatase from peripheral nerves. Biochimica et Biophysica Acta, 23: 394–401.

Tavalin, S. J., Ellis, E. F., Satin, L. S. (1997). Inhibition of the electrogenic Na pump underlies delayed depolarization of cortical neurons after mechanical injury or glutamate. J. Neurophysiol, 77: 632–8.

Thorpe G. W., Fong C. S., Alic N., Higgins V. J., Dawes I. W. (2004). Cells have distinct mechanisms to maintain protection against different reactive oxygen species: Oxidative-stress-response genes. Proc. Natl. Acad. Sci. U. S. A 101(17): 6564–6569

Toledano, M. B., Delaunay, A., Biteau, B., Spector, D., Azevedo, D. (2003). Oxidative stress responses in yeast. In Yeast Stress Responses Edited by: Hohman S, Mager WH. Berlin: Springer-Verlag: 305–387.

Toyoshima C., Kanai R., Cornelius F. (2011). First crystal structures of Na+,K+-ATPase: New light on the oldest ion pump. Structure, 19: 1732–1738.

Wong H. L, Liebler D. C. (2008) Mitochondrial protein targets of thiol-reactive electrophiles. Chem. Res. Toxicol, 21:796–804.

Zahler, R., Zhang, Z. T., Manor, M., Boron, W. F. (1997). Sodium kinetics of Na,K-ATPase alpha isoforms in intact transfected HeLa cells. J. Gen. Physiol, 110: 201–13.

